# DeepHeme: A generalizable, bone marrow classifier with hematopathologist-level performance

**DOI:** 10.1101/2023.02.20.528987

**Authors:** Gregory M. Goldgof, Shenghuan Sun, Jacob Van Cleave, Linlin Wang, Fabienne Lucas, Laura Brown, Jacob D. Spector, Leonardo Boiocchi, Jeeyeon Baik, Menglei Zhu, Orly Ardon, Chuanyi M. Lu, Ahmet Dogan, Dmitry B. Goldgof, Iain Carmichael, Sonam Prakash, Atul J. Butte

**Affiliations:** Department of Laboratory Medicine, University of California, San Francisco, CA, USA; Bakar Computational Health Sciences Institute, University of California, San Francisco, CA, USA; Department of Pathology and Laboratory Medicine, Memorial Sloan Kettering Cancer Center, New York, NY, USA; Department of Pathology, Brigham and Women’s Hospital/Harvard Medical School, Boston, MA, USA; Department of Laboratory Medicine, Boston Children’s Hospital/Harvard Medical School, Boston, MA, USA; Department of Laboratory Medicine, Veterans Affairs Medical Center, San Francisco, CA, USA; Department of Computer Science, University of South Florida, Tampa, FL, USA; Department of Statistics, University of California, Berkeley, CA, USA

**Author notes:** these authors contributed equally to this work (Gregory M. Goldgof, Shenghuan Sun).

## Abstract

Morphology-based classification of cells in the bone marrow aspirate (BMA) is a key step in the diagnosis and management of hematologic malignancies. However, it is time-intensive and must be performed by expert hematopathologists and laboratory professionals. We curated a large, high-quality dataset of 41,595 hematopathologist consensus-annotated single-cell images extracted from BMA whole slide images (WSIs) containing 23 morphologic classes from the clinical archives of the University of California, San Francisco. We trained a convolutional neural network, DeepHeme, to classify images in this dataset, achieving a mean area under the curve (AUC) of 0.99. DeepHeme was then externally validated on WSIs from Memorial Sloan Kettering Cancer Center, with a similar AUC of 0.98, demonstrating robust generalization. When compared to individual hematopathologists from three different top academic medical centers, the algorithm outperformed all three. Finally, DeepHeme reliably identified cell states such as mitosis, paving the way for image-based quantification of mitotic index in a cell-specific manner, which may have important clinical applications.

## Introduction

Hematologic malignancies such as leukemias, lymphomas, myelodysplastic syndromes, plasma cell neoplasms, and their precursor states, represent roughly 10% of cancer cases and cancer deaths worldwide^1^. Additionally, there are many non-neoplastic hematologic disorders originating in the bone marrow, from sickle cell disease, to iron deficiency anemia, to bone marrow failure syndromes^2^. For all these diseases, bone marrow aspirates (BMAs) are a central diagnostic tool. They allow assessment of morphology and relative distribution of cell types, which guides diagnostic and treatment decisions. Importantly, the exact quantification of cell subsets such as blasts, blast-equivalents or plasma cells is required to assign distinct diagnostic categories, with significant implications for treatment and prognosis^3–8^.

Despite its frequent clinical use, this process is technically challenging and time-intensive. Common sources of heterogeneity include different sample processing and staining procedures, overlapping morphologic features, and interobserver variability^9^. The limited number of professionals with these skills must balance the time needed to make an accurate diagnosis, with the clinical need for a fast and accurate diagnosis. For diseases such as acute promyelocytic leukemia, treatment delays can lead to death within days^10,11^. However, accuracy and reproducibility are just as important as speed, since diagnostic ambiguity and interobserver variability may have severe clinical implications. For example, when distinguishing acute myeloid leukemia with monocytic differentiation from chronic myelomonocytic leukemia, precise quantification of blast-equivalents from mature monocytes can be extremely challenging, lead to overestimation or underestimation of blast percentage, and consequently, overtreatment or undertreatment of patients^12–17^.

Morphologic examination of BMAs has been a major bottleneck in the adoption of WSI systems in hematopathology clinical practice. While whole slide scanners are becoming widely used for paraffin-embedded tissue specimens, BMAs continue to be examined under microscopes. Cell counting and classification done this way is both time consuming and error-prone, as it is possible to miss fields of view when using a microscope^9^. The ability to perform accurate classification of BMA cells using a comparable WSI image would eliminate this bottleneck and pave the way for digitization of these services and the development of algorithms to automate workflows and provide diagnostic support.

Morphologic examination of BMAs has been a major bottleneck to adoption of WSI systems in hematopathology clinical practice. While whole slide scanners are becoming widely adopted for paraffin-embedded tissue specimens, BMAs continue to be examined under microscopes, using a range of magnification (400x-1000x) and either light or oil microscopy. Cell counting and classification done this way is both time consuming and error prone, as it is very possible to miss fields of view under this level of magnification^9^. The ability to perform accurate classification of BMA cells using a comparable WSI image would eliminate this bottleneck to hematopathology clinical practice. Eliminating this bottleneck would lead to the digitization of these services and allow for the deployment of algorithms that help automate the current workflow, as well as future algorithms developed using this digitized data that can improve the quality of diagnoses.

## Results

### Case Identification, Whole Slide Imaging, and Image Annotation

50 aspirate slides from 50 unique patients with normal BMA morphology were selected from the UCSF Parnassus adult hospital and UCSF Benioff Children’s Hospital between 2017 and 2020. These BMAs were performed for diagnosis, staging, and monitoring and showed uninvolved marrow and normal hematopoiesis. Patient’s ages ranged from six months to 79 years, with 16% of cases originating from the pediatric hematopathology service. The gender composition was roughly equal (48% male, 52% female). Race/ethnicity was as follows: White: 78%, Hispanic: 12%, Asian 8%, and Black 2%. No other patient information was used or kept with the slide images, which were de-identified during the scanning process. WSIs were created using 400x-equivalent magnification.

The UCSF slide set was separated at this stage into 40 slides used for training and validation. 10 slides were kept as a hold-out test set to ensure accurate reporting of the model’s performance on unseen patient cases. A library of 41,595 images was assigned into one of 23 classes by consensus decision of an expert panel of three hematopathologists (Table 1). 30,394 images in the training set and 8,507 images in the test set.

**Table 1.**
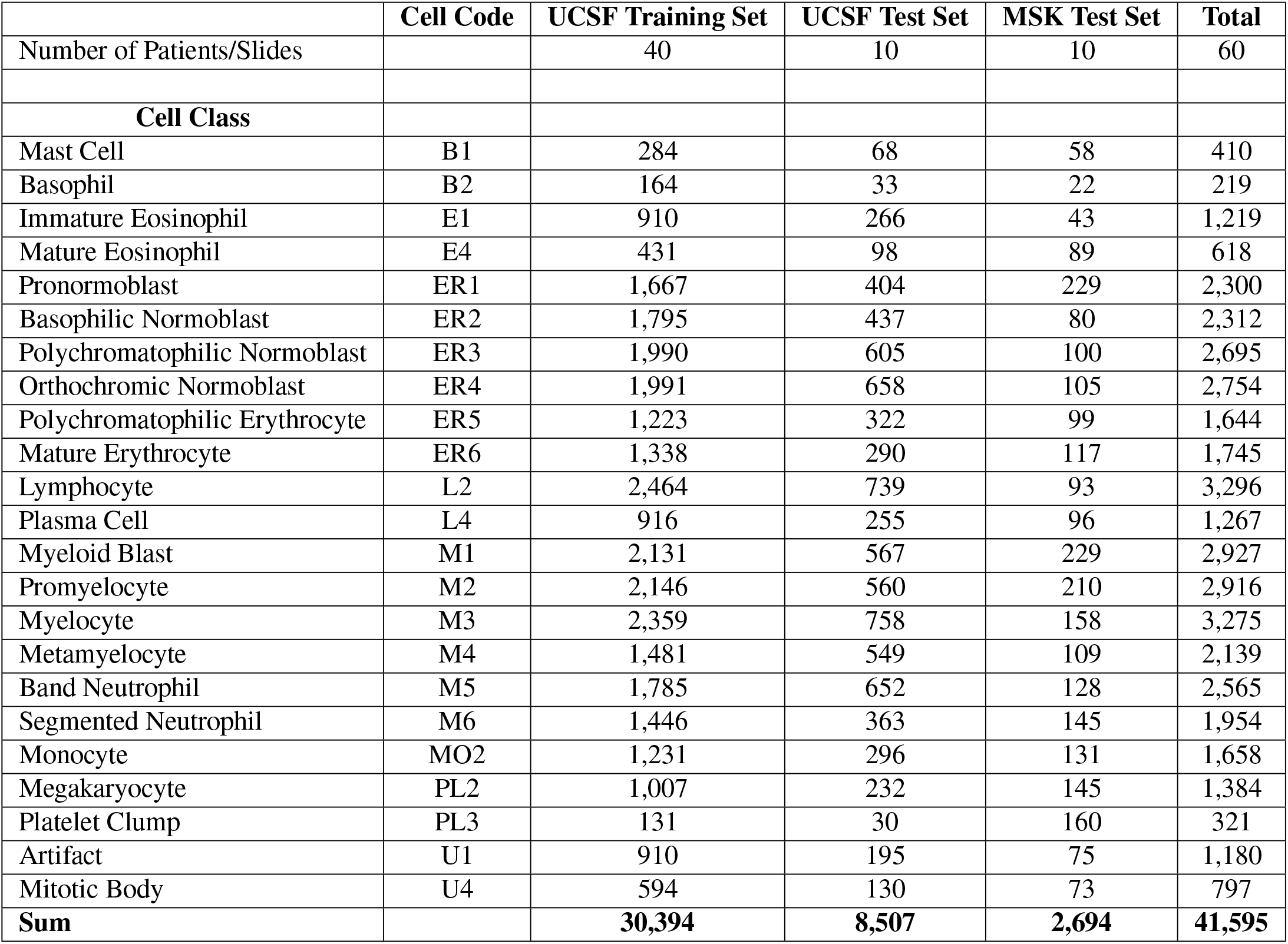
Datasets. This table shows the Number of hematopathologist-labeled, single cell images, per cell category, evaluated in the training, test, and external validation sets. Training and test sets were separated at the slide level to avoid testing on images from slides on which training had been performed. Each slide was obtained from a unique patient undergoing bone marrow evaluation, with results showing normal hematopoiesis.

|  | Cell Code | UCSF Training Set | UCSF Test Set | MSK Test Set | Total |
| --- | --- | --- | --- | --- | --- |
| Number of Patients/Slides |  | 40 | 10 | 10 | 60 |
| **Cell Class** |  |  |  |  |  |
| Mast Cell | B1 | 284 | 68 | 58 | 410 |
| Basophil | B2 | 164 | 33 | 22 | 219 |
| Immature Eosinophil | E1 | 910 | 266 | 43 | 1,219 |
| Mature Eosinophil | E4 | 431 | 98 | 89 | 618 |
| Pronormoblast | ER1 | 1,667 | 404 | 229 | 2,300 |
| Basophilic Normoblast | ER2 | 1,795 | 437 | 80 | 2,312 |
| Polychromatophilic Normoblast | ER3 | 1,990 | 605 | 100 | 2,695 |
| Orthochromic Normoblast | ER4 | 1,991 | 658 | 105 | 2,754 |
| Polychromatophilic Erythrocyte | ER5 | 1,223 | 322 | 99 | 1,644 |
| Mature Erythrocyte | ER6 | 1,338 | 290 | 117 | 1,745 |
| Lymphocyte | L2 | 2,464 | 739 | 93 | 3,296 |
| Plasma Cell | L4 | 916 | 255 | 96 | 1,267 |
| Myeloid Blast | M1 | 2,131 | 567 | 229 | 2,927 |
| Promyelocyte | M2 | 2,146 | 560 | 210 | 2,916 |
| Myelocyte | M3 | 2,359 | 758 | 158 | 3,275 |
| Metamyelocyte | M4 | 1,481 | 549 | 109 | 2,139 |
| Band Neutrophil | M5 | 1,785 | 652 | 128 | 2,565 |
| Segmented Neutrophil | M6 | 1,446 | 363 | 145 | 1,954 |
| Monocyte | MO2 | 1,231 | 296 | 131 | 1,658 |
| Megakaryocyte | PL2 | 1,007 | 232 | 145 | 1,384 |
| Platelet Clump | PL3 | 131 | 30 | 160 | 321 |
| Artifact | U1 | 910 | 195 | 75 | 1,180 |
| Mitotic Body | U4 | 594 | 130 | 73 | 797 |
| **Sum** |  | **30,394** | **8,507** | **2,694** | **41,595** |

The 23 image classes represent all cell types included in a standard bone marrow differential, as well as differentiation stages of trilineage hematopoietic cells (Figure 1b). The full spectrum of erythroid and neutrophil maturation was included, from proerythroblast to mature erythrocyte and from myeloid blast to segmented neutrophil, respectively. Along the megakaryocytic lineage, megakaryocytes and platelet clumps were assessed. The lymphoid lineage included lymphocytes and plasma cells. Eosinophils were separated into mature eosinophils with segmented nuclei and immature eosinophils. In addition, the set included monocytes, basophils and mast cells.

**Figure 1.**
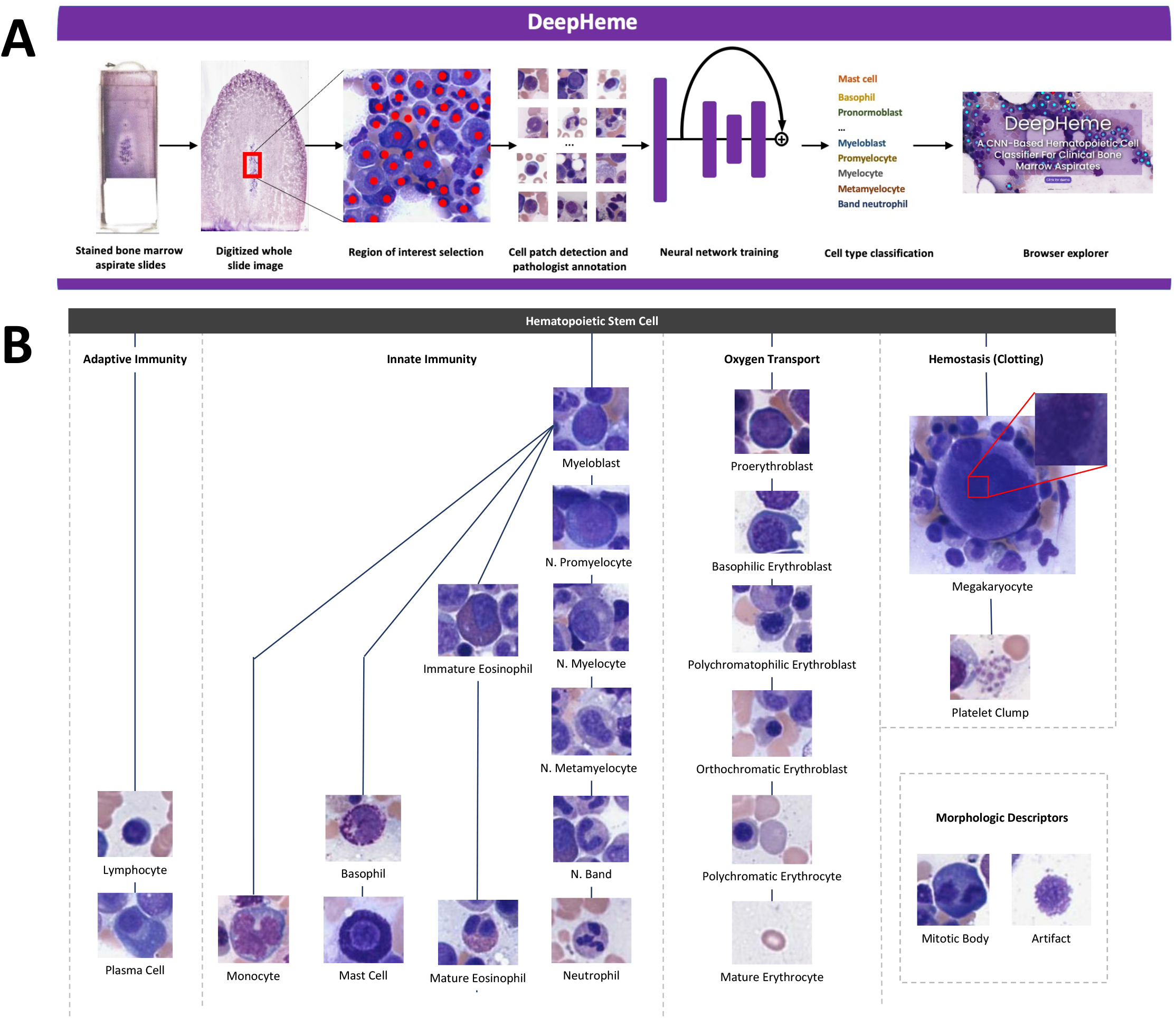
**A) Experimental workflow**. Whole slide images of bone marrow aspirates were digitized using whole slide scanners. Regions of interest were selected by hematopathologists and the location and classification of cells was labeled by a consensus of three hematopathologists. The single cell images were used to train and test convolutional neural networks with ResNext-50 architecture to produce DeepHeme, an algorithm that classifies single cell images into 23 different cell classes. A web application was built, where scientists can interact with the DeepHeme algorithm (https://www.hemepath.ai/deepheme.html). **B) Cell classes, lineage trajectories, and physiologic functions**. A diagram of cells and morphologic labels included in the study, as well as their relationship to each other in the hematopoietic tree. In addition to classes of cells, two important morphologic categories were included: mitotic body and artifact.

Additional classes include artifacts and mitotic bodies to probe for cellular states. The presence of mitotic boides is a proxy for the mitotic rate of the sample, which is itself a clinically prognostic biomarker^18–21^. Because of the differences in the relative distribution of bone marrow cell types, special efforts were made to identify additional examples of rare classes including myeloid blasts, basophils, mast cells, and mitotic bodies^22^.

### Neural Network Structure, Training and Testing

We used the ResNeXt-50 architecture^23^, which showed good performance in classifying bone marrow cells by Matek et al^24^. ResNeXt is a convolutional neural network architecture, which combines elements of of VGG^25^, ResNet^26^ and Inception models^27^. The network was initialized using weights from ImageNet-9 and then trained using bone marrow cell images^28^.

All separations of training and test sets were performed at the slide/ patient level to avoid repeat test evaluation on images from slides that had been used in training, even if the individual image instance was different. The 30,394 images from the 40 slides in the training set were split into training and validation sets using an 80:20 split and 5-fold cross validation. Testing was performed on the 8,507 images from the 10 slide test set.

In order to counteract an imbalanced distribution of cell types, up-sampling was used to equilibrate the classes prior to data augmentation, which resulted in 50,000 images per class. Shape augmentation included rotations, vertical and horizontal flips, shears and resize. Color augmentation included contrast, brightness, Gaussian noise and stain-color transformations^29,30^.

### Bone Marrow Cell Classification Performance

All performance metrics were averaged across the five networks created as a result of 5-fold cross validation process. Performance of the DeepHeme model was evaluated using AUROC, F1 score, accuracy„ precision, and recall (sensitivity) (Table 2, Supplemental Table 1). The mean AUC, precision, and recall across all classes was 0.99, 0.89, and 0.89, respectively. We define high performance as an average precision and recall both above 0.8. Using this metric, 19 of the 23 classes of cells achieved high performance. The algorithm achieved an AUC above 0.98 on all classes, except basophil (AUC=0.94), which also happened to have the smallest number of training images.

**Table 2.**
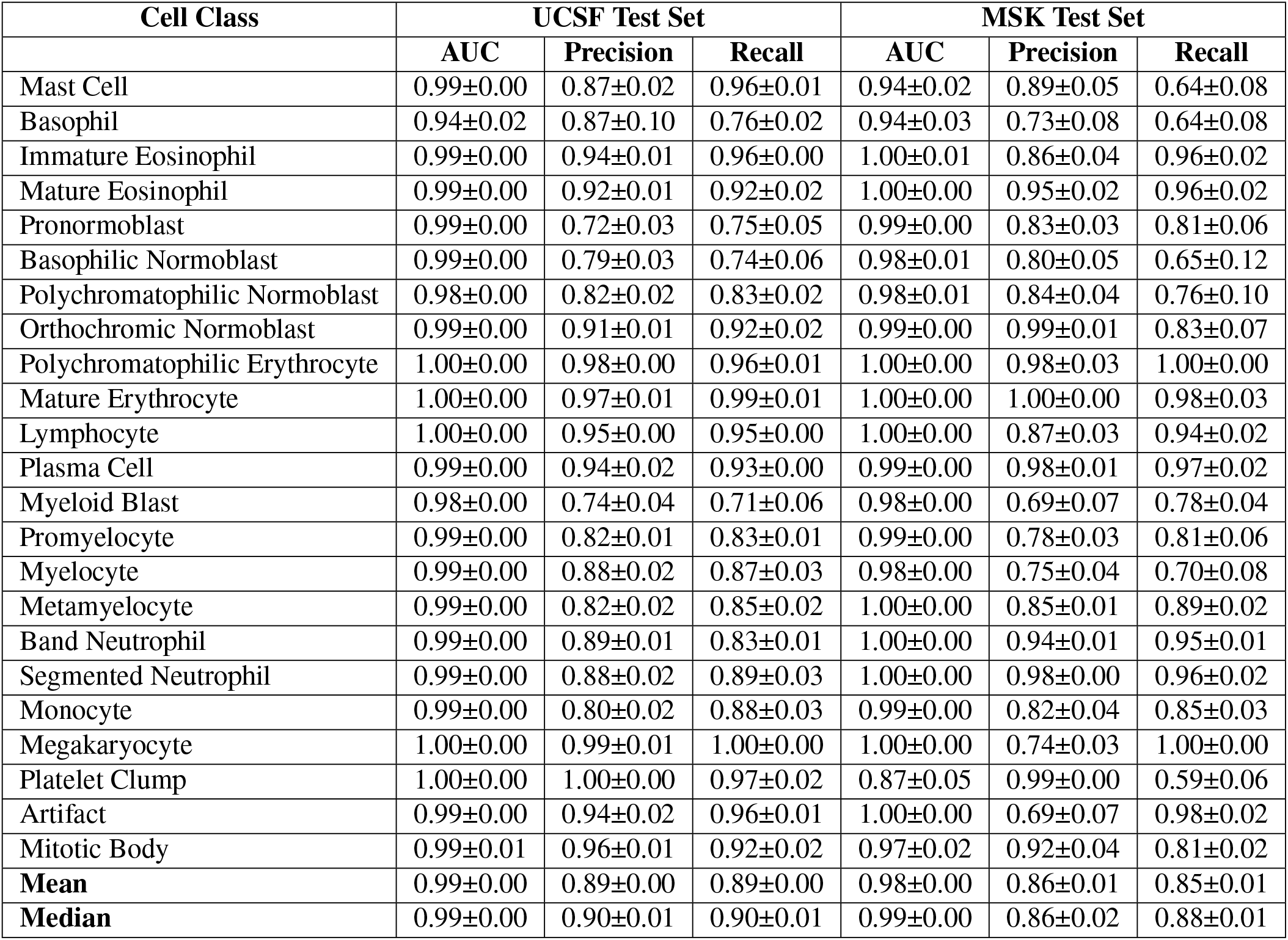
DeepHeme performance and external validation metrics. AUC, precision and recall were calculated for the hold-out UCSF test set and the external MSK test set. The mean AUC, precision, and recall across all classes for the UCSF test set were 0.99, 0.89 and 0.89, respectively. There were 1%, 3%, and 4% decreases for mean AUC, precision, and recall between the datasets, demonstrating the model’s high level of generalizability. F1-score and accuracy are available in the Supplemental Tables 1 and 2.

| Cell Class | UCSF Test Set |  |  | MSK Test Set |  |  |
| --- | --- | --- | --- | --- | --- | --- |
|  | AUC | Precision | Recall | AUC | Precision | Recall |
| Mast Cell | 0.99±0.00 | 0.87±0.02 | 0.96±0.01 | 0.94±0.02 | 0.89±0.05 | 0.64±0.08 |
| Basophil | 0.94±0.02 | 0.87±0.10 | 0.76±0.02 | 0.94±0.03 | 0.73±0.08 | 0.64±0.08 |
| Immature Eosinophil | 0.99±0.00 | 0.94±0.01 | 0.96±0.00 | 1.00±0.01 | 0.86±0.04 | 0.96±0.02 |
| Mature Eosinophil | 0.99±0.00 | 0.92±0.01 | 0.92±0.02 | 1.00±0.00 | 0.95±0.02 | 0.96±0.02 |
| Pronormoblast | 0.99±0.00 | 0.72±0.03 | 0.75±0.05 | 0.99±0.00 | 0.83±0.03 | 0.81±0.06 |
| Basophilic Normoblast | 0.99±0.00 | 0.79±0.03 | 0.74±0.06 | 0.98±0.01 | 0.80±0.05 | 0.65±0.12 |
| Polychromatophilic Normoblast | 0.98±0.00 | 0.82±0.02 | 0.83±0.02 | 0.98±0.01 | 0.84±0.04 | 0.76±0.10 |
| Orthochromic Normoblast | 0.99±0.00 | 0.91±0.01 | 0.92±0.02 | 0.99±0.00 | 0.99±0.01 | 0.83±0.07 |
| Polychromatophilic Erythrocyte | 1.00±0.00 | 0.98±0.00 | 0.96±0.01 | 1.00±0.00 | 0.98±0.03 | 1.00±0.00 |
| Mature Erythrocyte | 1.00±0.00 | 0.97±0.01 | 0.99±0.01 | 1.00±0.00 | 1.00±0.00 | 0.98±0.03 |
| Lymphocyte | 1.00±0.00 | 0.95±0.00 | 0.95±0.00 | 1.00±0.00 | 0.87±0.03 | 0.94±0.02 |
| Plasma Cell | 0.99±0.00 | 0.94±0.02 | 0.93±0.00 | 0.99±0.00 | 0.98±0.01 | 0.97±0.02 |
| Myeloid Blast | 0.98±0.00 | 0.74±0.04 | 0.71±0.06 | 0.98±0.00 | 0.69±0.07 | 0.78±0.04 |
| Promyelocyte | 0.99±0.00 | 0.82±0.01 | 0.83±0.01 | 0.99±0.00 | 0.78±0.03 | 0.81±0.06 |
| Myelocyte | 0.99±0.00 | 0.88±0.02 | 0.87±0.03 | 0.98±0.00 | 0.75±0.04 | 0.70±0.08 |
| Metamyelocyte | 0.99±0.00 | 0.82±0.02 | 0.85±0.02 | 1.00±0.00 | 0.85±0.01 | 0.89±0.02 |
| Band Neutrophil | 0.99±0.00 | 0.89±0.01 | 0.83±0.01 | 1.00±0.00 | 0.94±0.01 | 0.95±0.01 |
| Segmented Neutrophil | 0.99±0.00 | 0.88±0.02 | 0.89±0.03 | 1.00±0.00 | 0.98±0.00 | 0.96±0.02 |
| Monocyte | 0.99±0.00 | 0.80±0.02 | 0.88±0.03 | 0.99±0.00 | 0.82±0.04 | 0.85±0.03 |
| Megakaryocyte | 1.00±0.00 | 0.99±0.01 | 1.00±0.00 | 1.00±0.00 | 0.74±0.03 | 1.00±0.00 |
| Platelet Clump | 1.00±0.00 | 1.00±0.00 | 0.97±0.02 | 0.87±0.05 | 0.99±0.00 | 0.59±0.06 |
| Artifact | 0.99±0.00 | 0.94±0.02 | 0.96±0.01 | 1.00±0.00 | 0.69±0.07 | 0.98±0.02 |
| Mitotic Body | 0.99±0.01 | 0.96±0.01 | 0.92±0.02 | 0.97±0.02 | 0.92±0.04 | 0.81±0.02 |
| **Mean** | 0.99±0.00 | 0.89±0.00 | 0.89±0.00 | 0.98±0.00 | 0.86±0.01 | 0.85±0.01 |
| **Median** | 0.99±0.00 | 0.90±0.01 | 0.90±0.01 | 0.99±0.00 | 0.86±0.02 | 0.88±0.01 |

To better understand the severity of misclassification, we constructed a confusion matrix that plots true versus predicted labels (Supplemental Figure 1). Most miscategorizations occurred between developmentally and morphologically adjacent cell types. For example, most misclassified metamyelocytes (M3) were classified as either myelocytes (M2) or band neutrophils (M5), the precursor or successor cell stage in normal hematopoiesis. Since these categories are not distinct groups, but rather classifications on a continuous biologic spectrum, we would expect this result based on ambiguous cases, even among human experts.

The most common misclassification was between myeloid and erythroid blasts, with 14% of myeloid blasts misclassified as erythroid blasts and 12% of erythroid blasts misclassified as myeloid blasts. These classes are developmentally adjancent and very similar in morphology. No previous work has attempted to distinguish them computationally at 400x-equivalent resolution, although Choi et al. did so when using 1000x oil microscopy^31^.

### Model Learns Underlying Hematopoietic Developmental Relationships

Uniform Manifold Approximation and Projection for Dimension Reduction^32^ (UMAP) was used to create a two-dimensional representation of the high-dimensional features learned by DeepHeme. This was done to visualize and explore how the different classes are being grouped together or separated from each other(Figure 2a). Towards the top left of the UMAP, we see that myeloid blast (myeloblast) and proerythroblast are connected by a thick bridge of cells, representing the theoretical hematopoietic stem cell population from which they both differentiate. Cascading bridges downward, and to the right, represent neutrophil and erythroid development, respectively (Figure 2b and 2c).

**Figure 2.**
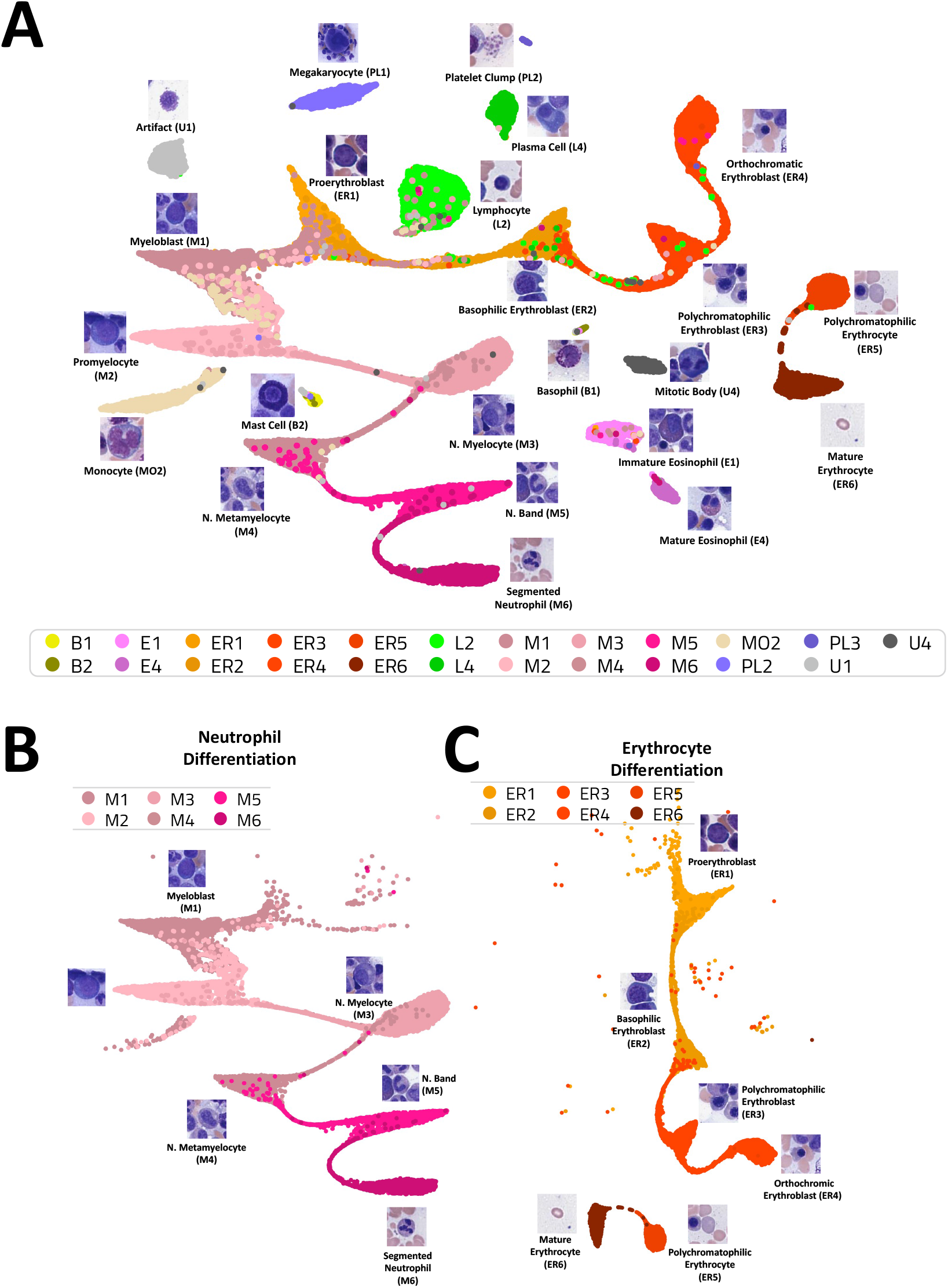
**A) UMAP embedding of extracted features recapitulates biological relationships**. The shape of the UMAP recapitulates major aspects of hematopoiesis, of which the untrained neural network has no prior knowledge, suggesting it has been learned from the training images. Bridges between clusters reflect the natural continuum and lineage trajectories between adjacent cell types. **B) Neutrophil differentiation**. The complete spectrum of neutrophil development from myeloblast to segmented neutrophil has been learned by the algorithm. **C) Erythrocyte differentiation**. Similarly, the full spectrum of erythroid development has also been inferred by the algorithm. The break between orthochromatic erythroblasts (ER5) and polychromatic erythrocytes (ER6) likely reflects their clear morphologic boundary (the presence of any nuclear remnants). Such clear morphologic boundaries do not exist between other cell categories, which are defined based on multiple, subjective features.

The only clear break in the erythroid lineage is between orthochromic erythroblasts and polychromatic erythrocytes. These are also the only two cell classes that have a clearly defined morphologic division that separates them (the presence or absence of a nucleus). For the rest of neutrophil and erythrocyte development, the dividing line between adjacent cell classes requires interpretation based on multiple subjective parameters.

In fact, a bridge is seen between nearly all directly related hematopoietic cell types. Their is a notable absence of a bridge from the hematopoietic stem cell to lymphocytes or plasma cells, as they do not undergo development in the marrow, but rather return to the marrow after maturation in the thymus, lymph nodes, and other lymphoid tissues. The inclusion of intermediate cell types such as hematogones or lymphoplasmacytoid cells in the future may link these groups. Other un-bridged, related cells are likely missing intermediate cell types due to their infrequency in the marrow.

The algorithm also learned morphologic relationships across lineages. For example, all five cell classes in our dataset with high nuclear-to-cytoplasmic (N:C) ratio co-localize, while only cell clusters that are directly related to each other are attached by bridges (Supplemental Figure 2). Of note, the unconnected class in this group, lymphocytes, is also the only mature cell type. These findings suggest N:C ratio can be inferred by distance from this cluster, which is a morphologic feature common to many malignancies, and maturity can be inferred by proximity to, and interconnection with, the myeloblast and proerythroblast cell clusters.

Another example is the positioning of the mitotic bodies cluster. The mitotic bodies in this study broadly derive from three morphologic groups based on cytoplasmic color, which correspond to the lineage of the cell undergoing mitosis. Mitotic cells in the neutrophil lineage have a pale pink cytoplasm, those in the eosinophilic lineage contain bright pink cytoplasm and those in the erythroid lineage have deep blue cytoplasm. The mitotic body cluster is placed by the UMAP directly between these three lineage cluster sets, perhaps reflecting the relationship of the mitotic body class to these three lineages. These findings suggest further subclassification of mitotic bodies by lineage is possible based on cytoplasm color.

This UMAP has recapitulated much of the hematopoietic structure known to biologists as a result of decades of research (Figure 1b). It is important to point out that the deep learning algorithm was not given any information about the underlying relationships between the classes. Rather, it has reconstructed elements of the hematopoietic tree based on morphology alone. The fact that UMAP recapitulates biological relationships suggests that the algorithm is learning relevant morphologic features rather than confounders or shortcuts, which would not recapitulate biological relationships. These findings also suggest the possibility of identifying novel biological relationships by combining single cell image datasets with dimension reduction methods.

### Comparison with Clinical Experts

To determine whether or not DeepHeme achieves clinical level accuracy, we compared its results to pathologists’ assessments. We selected three subspecialty hematopathologists from three different well-established cancer centers (MSKCC, UCSF and Brigham and Women’s Hospital) and asked them to perform the same classification task as the algorithm. Specialists were blinded to each other, unlike the gold-standard labels, which were defined by consensus. Twenty-five images from each of the 23 classes from the UCSF test set were randomly chosen for review. Figure 3a summarizes the performance of the reviewers versus AI. DeepHeme achieved hematopathologist-level or better performance on all 23 classes, with a mean precision and recall of (0.90±0.00, 0.90±0.00) versus (0.78±0.05, 0.76±0.06) for hematopathologists (Figure 3a and Supplemental Tables 3-5). In terms of variation, the mean standard deviation for precision and recall across the 23 classes was (0.03±0.02, 0.04±0.02) for the AI, versus (0.08±0.06, 0.10±0.00.05) for hematopathologists. In terms of speed, the three hematopathologists averaged roughly 3 hours for the task of labeling 575 images. By contrast, DeepHeme performed the same task in 0.36 seconds, or approximately 30,000 times faster than the experts.

**Figure 3.**
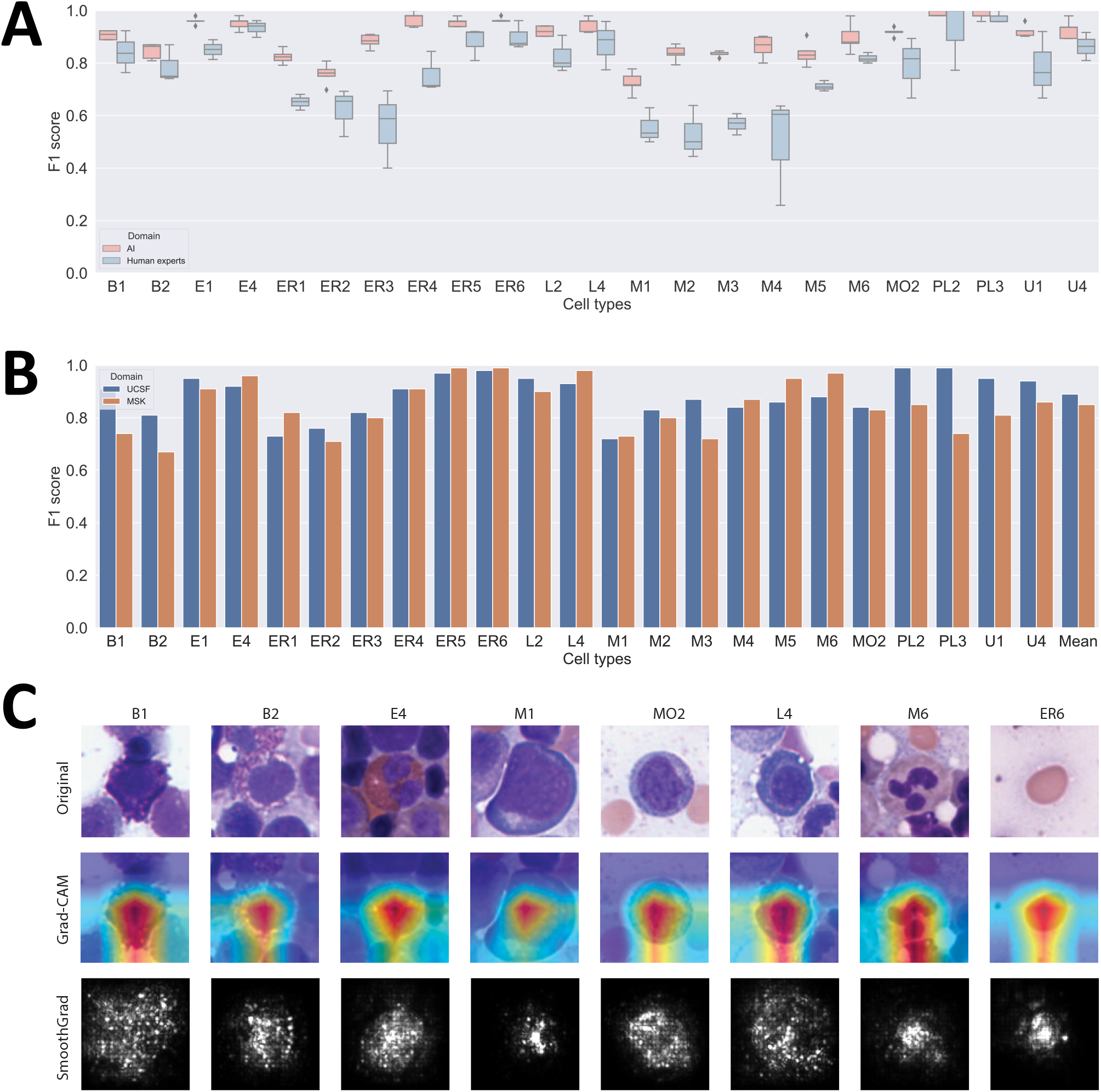
**A) Comparison of DeepHeme algorithm to expert hematopathologist evaluation**. Each cell was first categorized by three independent hematopathologists blinded to each other. 25 images from each of the 23 classes from the UCSF test set were randomly chosen to be re-labelled by three individual hematopathologists from three different institutions to assess the performance limits of experts, as well as variability among them (green). DeepHeme (red) achieved hematopathologist-level performance or superiority in all 23 classes. **B) Model Generalizability**. Comparison of performance of DeepHeme on the UCSF test set, versus the external MSK set. We see a modest variation in performance across most cell classes, with a mean F1 score dropping from 0.89±0.00 to 0.85±0.01. **C) Saliency Maps**. This figure shows randomly selected saliency maps from 8 classes using different mapping algorithms. Row 1: Original. Row 2: Grad-CAM. Row 3: SmoothGrad. In each case, the map demonstrates that DeepHeme focuses on the cell of interest rather than adjacent cells or non-cellular components of image.

### Model Generalizability

Multi-site generalization has proven difficult to achieve in many areas of medical computer vision^33,34^. While no algorithm has demonstrated multi-site generalization for this problem, it is a necessary precondition to widespread adoption, since building pathologist-labelled training sets at each individual hospital is unreasonable. To evaluate DeepHeme’s ability to generalize to an external dataset, we next tested the classifier on images from a completely independent hospital system, Memorial Sloan Kettering Cancer Center (MSK). The MSK dataset includes 2,694 images from 10 randomly selected de-identified normal slides from the hematopathology service (Table 1). Images were scanned using a Leica Aperio AT2, de-identified, and then sent to UCSF for annotation. The same annotation strategy was used for the external dataset, to ensure that differences in performance were not the consequence of inter-annotator variability. Table 2 and Figure 3b summarize the performance of DeepHeme on the external dataset. We see a decrease of only 1%, 3%, and 4%, for mean AUC, precision, and recall, respectively between the UCSF and MSK datasets (Table 2), demonstrating robust generalization. The largest performance drop was in PL3 (platelet clumps). Hematopathologist review of the dataset revealed that platelet clumps in the MSK dataset were larger and more numerous due to variation in slide preparation (laboratory artifacts).

### Saliency Mapping

Saliency refers to what is noticeable or important in an image. The saliency map for an image in the context of convolution neural networks represents the significant pixels in the picture that affect the class score of the network’s prediction. Saliency maps were used to visualize which parts of the image were most important to the classification. If the algorithm were focusing outside the cell of interest, it would demonstrate that its performance is based on confounding factors, rather than medically-relevant image features. This type of error, called shortcut learning, is common in deep learning models^33,34^. Figure 3c shows eight randomly selected images from different classes, as well as their saliency maps using different mapping algorithms^25,35–37^. Using this approach, we see that DeepHeme is correctly focusing on the cell of interest and not adjacent cells or non-cellular components of the image.

### Comparison with Other Bone Marrow Cell Classifiers

Direct comparison with shared test sets from other published studies in this area is not possible since there is significant variation in class definitions and limited data availability. However, Table 3 compares DeepHeme to recent high-quality publications in this field. Chandradevan et al. was the first study to perform analysis using deep learning on bone marrow slides from WSIs, a breakthrough at the time. This study is limited by a relatively small dataset, limited image classes, and limited reporting on performance13^38^. Aside from DeepHeme, it is the only other paper to employ a consensus of multiple hematopathologists to develop its labels.

**Table 3.** Comparison to other deep-learning-based bone marrow cell classifiers. This table compares DeepHeme to other works that use CNNs to classify single cell images from 400x-equivalent images. All image totals are for the entire study, including training and test sets. ‘High performance’ is defined as >0.8 precision and recall. ‘Images in smallest class’ is a measure of class imbalance and is included because tiny training sets for individual classes generally yields low performance for those classes. DeepHeme matches or outperforms currently published algorithms across all metrics except total number of images used in the study.

|  | Chandradevan et al. | Lewis et al. | Matek et al. | Tayebi et al. | DeepHeme |
| --- | --- | --- | --- | --- | --- |
| **Annotation Standard** | Hematopathologist consensus | Not reported | Clinical Laboratory Staff | Hematopathologist | Hematopathologist consensus |
| **Image Source** | WSI (light) | WSI (light) | Microscope Camera under Oil | WSI (light) | WSI (light) |
| **Images** | 9,269 | 23,609 | 171,374 | 26,782 | 41,595 |
| **Image Classes** | 12 | 16 | 21 | 19 | 23 |
| **Images in smallest class** | 62 | 155 | 8 | 7 | 219 |
| **Classes with high performance** | Not reported | Not reported | 7 | 6 | 19 |
| **Mean F1-Score** | Not reported | Not reported | 0.67 | 0.78 | 0.89 |
| **Mean Precision** | Not reported | Not reported | 0.61 | 0.83 | 0.89 |
| **Mean Recall** | Not reported | Not reported | 0.76 | 0.75 | 0.89 |
| **Slide Level External Validation** | No | Yes | No | Yes | Yes |
| **Expert Comparison** | No | No | No | No | Yes |
| **Institution Level External Validation** | No | No | No | No | Yes |

Matek et al. made significant improvements through the use of a large-scale dataset and higher number of classes, however the dataset was not annotated by hematopathologists, which is the clinical gold-standard^24^. Both Matek and Chandradevan et al also appear to have separated their training and test sets once all images were cropped, meaning that their algorithms were tested on the same slides on which they were trained, likely leading to performance overestimates, overfitting and generalization issues. Matek et al. also used images taken using microscope cameras under oil at 400x, which may not generalize well to light-microscopy images and WSIs.

Lewis et al. and Tayebi et al. were the only other papers we identified that separated training and testing at the slide-level, demonstrating slide-level generalizability^39,40^. Tayebi et al and Matek et al each were limited by extreme class imbalances, which led to weak performance on many image classes. Some classes were trained on as few as 7 images. Consequently, they achieved high performance, as defined by >0.8 in precision and recall, on only 7 of 19, and 8 of 21, image classes, respectively. By contrast, DeepHeme achieved high performance on 19 of 23 image classes and achieved a higher mean F1-score (0.89 vs 0.78 vs 0.67). DeepHeme was the only algorithm to demonstrate generalization to an external setting. It was also the only algorithm to demonstrate performance comparable or superior to practicing hematopathologists.

### Website

A web application has been built at https://hemepath.ai/deepheme.html for scientists to interact with the DeepHeme algorithm. The application allows users to test the algorithm on images from either the UCSF or MSK test sets. They can also upload their own images. Images should be either cropped from 400x-equivalent WSIs or images captured from microscope cameras at 400x. Alternatively, users can capture pictures using microscope-mounted phone cameras and upload those images.

## Discussion

Classification and quantification of individual cell types in BMAs is an essential part of how hematopathologists diagnose hematological disorders. This task is not only time consuming and requires a high level of skill, but also involves considerable inter-observer variability. Only 200-500 individual cells are routinely quantified in standard clinical practice, affecting accurate quantification and meaning diseased cells may be missed.

C Computational tools can overcome these shortcomings, but they need to meet the following requirements: they should perform the original task faster, and they should have at least at the same level of quality as the human expert. The DeepHeme algorithm meets these requirements. First, it performs the diagnostic task in a fraction of a second, thereby vastly surpassing the human expert. Second, DeepHeme is the first algorithm for bone marrow cell classification to achieve high performance (defined as >0.8 precision and recall) on 20 out of 23 classes. Finally, it is also the first algorithm to demonstrate robust generalization, making it widely applicable across institutions.

We note the degree of interobserver variability among human experts in our study. This might reflect differences in local practices, different levels of experience, or the fact that cells were assessed on a single-cell level, where internal comparison to other cells was not allowed, a practice that is often used by human pathologists to “calibrate”. This variation underscores a key advantage of AI-based systems, which can analyze a large number of cells with low variation, thereby producing more accurate and reproducible diagnostic and prognostic signatures.

In addition, more nuanced subclassifications can be made routinely and reproducibly. For example, hematopathologists do not routinely sub-classify all erythroid maturation stages. However, subtle changes in developing erythroid cells may translate into clinically meaningful parameters or biomarkers that could advance the understanding of conditions such as myelodysplastic or bone marrow failure syndromes. AI-based assessments of BMAs could therefore enable the discovery of novel diagnostic and prognostic signatures.

The mitotic rate is a surrogate for cellular proliferation and is usually quantified using Ki-67 immunohistochemical stains on paraffin-embedded tissue, which is delayed by one or more days relative to the BMA. Ki-67 is only assessed semi-quantitatively, due to issues with variation in staining and inter-observer interpretation^41,42^. AI-assisted BMA mitotic rate assessment would have numerous advantages over Ki-67 staining. It does not require paraffin embedding or immunohistochemical staining, it allows more accurate quantification and “screening” of a larger number of cells, and it would allow assessment of individual cell morphology. DeepHeme is the first algorithm to classify BMA cells undergoing mitosis, making it a promising tool to develop biomarkers based on morphologic features of cells undergoing mitosis and mitotic rates of BMA samples.

While developed in healthy bone marrows, future work will focus on developing image libraries that represent the spectrum of hematologic disorders. By training on more extensive image libraries, an AI may learn morphologies associated with underlying genetic alterations, thereby allowing prediction of molecular or cytogenetic results. For example, in AML, many genetically-defined subtypes have associated morphologic features, such as distinct nuclear or cytoplasmic features^**?**^. Abnormal cell morphology is also a hallmark of myeloproliferative and myelodysplastic neoplasms.

In clinical workflows, DeepHeme can be used for screening and flagging of abnormal specimens, thereby substantially reducing the burden on human experts. Importantly, such screening would classify thousands to millions of cells per slide, rather than the 200-500 that humans typically classify, leading to more accurate cell counts and ensuring that areas that may have disease are not missed. For smaller clinics and resource limited settings, deployment in combination with a microscope camera or cell phone mounted microscope are promising approaches currently in development. By achieving high speed and diagnostic-grade accuracy, DeepHeme can help standardize clinical practice by reducing the variation seen in human experts, and pave the way for biomarker discovery in hematologic malignancies.

## Data Availability

Cell images will be published upon acceptance via The Cancer Imaging Archive and at www.hemepath.ai/DeepHeme.html. The code used for this project is downloadable on GitHub at https://github.com/goldgoflab/DeepHeme_training.

## Acknowledgements

We thank the members of the UCSF and MSK Hematopathology Services, as well as the MSKCC Warren Alpert Center. This work was supported by the National Institutes of Health (G.M.G., NHLBI R38HL143581 and NHLBI K38HL159128). This work was additionally supported by the UCSF Department of Laboratory Medicine.

## Author Contributions

GMG designed the study. GMG and SS developed the AI, analyzed the data, and drafted the manuscript. GMG and JVC built the image labelling software and demo site. AJB supervised the study. SP, LW, LB and CML provided hematopathology oversight. JDS provided advise on slide scanning and software development. AJB, DBG, IC provided image analysis and statistical oversight. GMG identified and scanned all UCSF slides. OA, AD and JB identified and scanned slides from MSKCC. GMG, LW, SP, LB, FL, and LB annotated images. All authors contributed to manuscript review and editing.

## Competing Interests

AJB is a co-founder and consultant to Personalis and NuMedii; consultant to Mango Tree Corporation, and in the recent past, Samsung, 10x Genomics, Helix, Pathway Genomics, and Verinata (Illumina); has served on paid advisory panels or boards for Geisinger Health, Regenstrief Institute, Gerson Lehman Group, AlphaSights, Covance, Novartis, Genentech, and Merck, and Roche; is a shareholder in Personalis and NuMedii; is a minor shareholder in Apple, Meta (Facebook), Alphabet (Google), Microsoft, Amazon, Snap, 10x Genomics, Illumina, Regeneron, Sanofi, Pfizer, Royalty Pharma, Moderna, Sutro, Doximity, BioNtech, Invitae, Pacific Biosciences, Editas Medicine, Nuna Health, Assay Depot, and Vet24seven, and several other non-health related companies and mutual funds; and has received honoraria and travel reimbursement for invited talks from Johnson and Johnson, Roche, Genentech, Pfizer, Merck, Lilly, Takeda, Varian, Mars, Siemens, Optum, Abbott, Celgene, AstraZeneca, AbbVie, Westat, and many academic institutions, medical or disease specific foundations and associations, and health systems. AJB receives royalty payments through Stanford University, for several patents and other disclosures licensed to NuMedii and Personalis. AJB’s research has been funded by NIH, Peraton (as the prime on an NIH contract), Genentech, Johnson and Johnson, FDA, Robert Wood Johnson Foundation, Leon Lowenstein Foundation, Intervalien Foundation, Priscilla Chan and Mark Zuckerberg, the Barbara and Gerson Bakar Foundation, and in the recent past, the March of Dimes, Juvenile Diabetes Research Foundation, California Governor’s Office of Planning and Research, California Institute for Regenerative Medicine, L’Oreal, and Progenity. AD receives research support form Roche and Takeda, and has been a consultant for EUSA Pharma, Incyte and Eli Lilly. The authors have declared that none of these potential competing interests affected the research or outcomes in any way.

## Online Content

### Methods

#### Case Identification and Whole Slide Imaging

Two new datasets were created to develop and test the performance of our deep learning algorithm, one from UCSF and one from MSK. All slides were randomly selected from the clinical service based on normal morphology and adequate specimen. For the UCSF dataset, the slides were stained using a version of the Wright-Giemsa stain (Fisherbrand modified Wright-Giemsa Stain Pack) and were either prepared at the UCSF Parnassus hospital or the UCSF Benioff Children’s Hospital between 2017 and 2020. WSIs were scanned at 400x-equivalent magnification using either a Leica Aperio AT Turbo or Leica Aperio AT2 and saved as de-identified .svs files. All MSK slides were scanned using the same methods using a Leica Aperio AT2. All slides were scanned using a high density of focus points and a single z-plane. Images include a range of image quality reflecting variations in stain intensity, slide preparation, and slide age.

#### Image Library Annotation

Images were annotated using annotation software developed in-house. To compensate for variations in slide preparation and stain intensity, as well as to replicate features of a manual microscope, the software’s viewer permits modification of brightness, contrast, and zoom. To compensate for variations in slide preparation and stain intensity, as well as to replicate features of a manual microscope, the software’s viewer permits modification of brightness, contrast, and zoom. Images from both the UCSF and MSK datasets were annotated using a 3-step process. Initial image classification was performed by a single pathologist. A second audit was performed by a single pathologist to remove any errors made with the first round of classification. Finally, a panel of three hematopathologists reviewed the final sorted cell lists to provide a consensus label that was used as the gold-standard.

For each image, the annotated cell is in the center of the image, based on the whole cell, not the nucleus, except for the following classes. Since megakaryocytes are larger than the field of view, image centers were placed in multiple non-overlapping locations within the megakaryocyte to capture different fields of view. For cells undergoing mitosis, the centers may have been placed in either the center of the mitotic figure, the center of the cell, or both. For platelet clumps, the center of the object was placed in the middle of the clump. Images were exported as 96×96 pixel PNGs with a resolution of 72px/inch.

#### Neural Network Structure, Training and Testing

In order to counteract a imbalanced distribution of cell types, up-sampling was used to equilibrate the classes prior to data augmentation. 20 times augmentation transformations using the Albumentations python library were performed to augment the shape and color, which resulted in approximately 50,000 images per class with 1.15 million augmented images in total1^30^. Shape augmentation included rotations, vertical and horizontal flips, shears and resize. Color augmentation included contrast, brightness, and Gaussian noise. In addition, we also performed stain-color augmentation transformations. A histopathology (HE)-based color augmentation was performed using the Stainlib python package^43^.

We used the ResNeXt-50 architecture^23^, which showed good performance in classifying bone marrow cells by Matek et al^24^. The network was initialized using weights from ImageNet-9 and then trained using bone marrow cell images^28^. Specifically, we modified the network input to accept images of the size 96 × 96 pixels, and adjust the number of output nodes, yielding the 23 overall cell types of our annotation scheme. The model took individual cell patches as the input and generated class predictions. After applying the softmax function to the output to generate probability distributions, the highest probability determined the cell class prediction results. The model was trained with batch size 1024, with an initial learning rate 0.001, for 30 epochs. The learning rate decayed every 5 epochs. Early stopping was used to prevent overfitting. Binary cross-entropy loss was used as the loss function with one-hot encoded targets. We used the Adam Optimization method for updating the model weights while reducing training error. All training was performed on NVIDIA TITAN RTX graphics processing units, where training of the ResNeXt model took approximately 12 hours of computing time. For training and validation, we used 40 images from slides from the UCSF dataset, whereas the rest 10 slides were used as the UCSF test set. 5-fold cross validation was performed on the training/validation set. We then trained 5 different networks for 13 epochs, where each network used a different fold for validation and the remaining 4 folds for training. All model tuning and parameters adjustment were performed during training and validation. All the numbers reported in the paper come from analysis of the unseen test sets. Results were then averaged across the 5 different cross-validation networks.

#### Saliency Mapping

In our research, we employed Grad-CAM and SmoothGrad saliency mapping algorithms to better understand the categorization choices made by these algorithms^25,35–37^.

#### UMAP Interpretation

Uniform Manifold Approximation and Projection for Dimension Reduction (UMAP) was used to represent the information that the deep learning classifier learned^32^. We embedded the extracted features represented in the flattened final convolutional layer of the network (1000 dimensions) into 2 dimensions for each member of the data set using the UMAP algorithm. UMAP works by using by using nearest-neighbor-descent technique to identify the closest neighbors. The nearest neighbors that were previously found are then connected to create a graph^44^. The next stage for UMAP is to map the approximation manifold to a lower-dimensional space, in our case two dimensions, after learning it from the higher-dimensional environment. To perform these calculations we used the umap-learn package in Python^45^.

## Supplemental Figures and Tables

**Table 1.** UCSF Results. Expanded performance metrics of DeepHeme algorithm on the UCSF test set.

| Cell Class | Cell Code | AUC | F1-Score | Accuracy | Precision | Recall |
| --- | --- | --- | --- | --- | --- | --- |
| Mast Cell | B1 | 0.99±0.00 | 0.91±0.01 | 1.00±0.00 | 0.87±0.02 | 0.96±0.01 |
| Basophil | B2 | 0.94±0.02 | 0.81±0.05 | 1.00±0.00 | 0.87±0.10 | 0.76±0.02 |
| Immature Eosinophil | E1 | 0.99±0.00 | 0.95±0.00 | 1.00±0.00 | 0.94±0.01 | 0.96±0.00 |
| Mature Eosinophil | E4 | 0.99±0.00 | 0.92±0.01 | 1.00±0.00 | 0.92±0.01 | 0.92±0.02 |
| Pronormoblast | ER1 | 0.99±0.00 | 0.73±0.01 | 0.97±0.00 | 0.72±0.03 | 0.75±0.05 |
| Basophilic Normoblast | ER2 | 0.99±0.00 | 0.76±0.02 | 0.98±0.00 | 0.79±0.03 | 0.74±0.06 |
| Polychromatophilic Normoblast | ER3 | 0.98±0.00 | 0.82±0.00 | 0.97±0.00 | 0.82±0.02 | 0.83±0.02 |
| Orthochromic Normoblast | ER4 | 0.99±0.00 | 0.91±0.01 | 0.99±0.00 | 0.91±0.01 | 0.92±0.02 |
| Polychromatophilic Erythrocyte | ER5 | 1.00±0.00 | 0.97±0.00 | 1.00±0.00 | 0.98±0.00 | 0.96±0.01 |
| Mature Erythrocyte | ER6 | 1.00±0.00 | 0.98±0.00 | 1.00±0.00 | 0.97±0.01 | 0.99±0.01 |
| Lymphocyte | L2 | 1.00±0.00 | 0.95±0.00 | 0.99±0.00 | 0.95±0.00 | 0.95±0.00 |
| Plasma Cell | L4 | 0.99±0.00 | 0.93±0.01 | 1.00±0.00 | 0.94±0.02 | 0.93±0.00 |
| Myeloid Blast | M1 | 0.98±0.00 | 0.72±0.02 | 0.96±0.00 | 0.74±0.04 | 0.71±0.06 |
| Promyelocyte | M2 | 0.99±0.00 | 0.83±0.00 | 0.98±0.00 | 0.82±0.01 | 0.83±0.01 |
| Myelocyte | M3 | 0.99±0.00 | 0.87±0.01 | 0.98±0.00 | 0.88±0.02 | 0.87±0.03 |
| Metamyelocyte | M4 | 0.99±0.00 | 0.84±0.01 | 0.98±0.00 | 0.82±0.02 | 0.85±0.02 |
| Band Neutrophil | M5 | 0.99±0.00 | 0.86±0.00 | 0.98±0.00 | 0.89±0.01 | 0.83±0.01 |
| Segmented Neutrophil | M6 | 0.99±0.00 | 0.88±0.01 | 0.99±0.00 | 0.88±0.02 | 0.89±0.03 |
| Monocyte | MO2 | 0.99±0.00 | 0.84±0.01 | 0.99±0.00 | 0.80±0.02 | 0.88±0.03 |
| Megakaryocyte | PL2 | 1.00±0.00 | 0.99±0.00 | 1.00±0.00 | 0.99±0.01 | 1.00±0.00 |
| Platelet Clump | PL3 | 1.00±0.00 | 0.99±0.01 | 1.00±0.00 | 1.00±0.00 | 0.97±0.02 |
| Artifact | U1 | 0.99±0.00 | 0.95±0.01 | 1.00±0.00 | 0.94±0.02 | 0.96±0.01 |
| Mitotic Body | U4 | 0.99±0.01 | 0.94±0.01 | 1.00±0.00 | 0.96±0.01 | 0.92±0.02 |
| Mean |  | 0.99±0.00 | 0.89±0.00 | 0.99±0.00 | 0.89±0.00 | 0.89±0.00 |
| Medium |  | 0.99±0.00 | 0.91±0.00 | 0.99±0.00 | 0.90±0.01 | 0.90±0.01 |

**Table 2.** MSK Results. Expanded performance metrics of DeepHeme algorithm on the MSK test set.

| Cell Class | Cell Code | AUC | F1-Score | Accuracy | Precision | Recall |
| --- | --- | --- | --- | --- | --- | --- |
| Mast Cell | B1 | 0.94±0.02 | 0.74±0.07 | 0.99±0.00 | 0.89±0.05 | 0.64±0.08 |
| Basophil | B2 | 0.94±0.03 | 0.67±0.05 | 0.99±0.00 | 0.73±0.08 | 0.64±0.08 |
| Immature Eosinophil | E1 | 1.00±0.01 | 0.91±0.03 | 1.00±0.00 | 0.86±0.04 | 0.96±0.02 |
| Mature Eosinophil | E4 | 1.00±0.00 | 0.96±0.01 | 1.00±0.00 | 0.95±0.02 | 0.96±0.02 |
| Pronormoblast | ER1 | 0.99±0.00 | 0.82±0.02 | 0.97±0.00 | 0.83±0.03 | 0.81±0.06 |
| Basophilic Normoblast | ER2 | 0.98±0.01 | 0.71±0.06 | 0.98±0.00 | 0.80±0.05 | 0.65±0.12 |
| Polychromatophilic Normoblast | ER3 | 0.98±0.01 | 0.80±0.07 | 0.99±0.00 | 0.84±0.04 | 0.76±0.10 |
| Orthochromic Normoblast | ER4 | 0.99±0.00 | 0.91±0.05 | 0.99±0.00 | 0.99±0.01 | 0.83±0.07 |
| Polychromatophilic Erythrocyte | ER5 | 1.00±0.00 | 0.99±0.02 | 1.00±0.00 | 0.98±0.03 | 1.00±0.00 |
| Mature Erythrocyte | ER6 | 1.00±0.00 | 0.99±0.02 | 1.00±0.00 | 1.00±0.00 | 0.98±0.03 |
| Lymphocyte | L2 | 1.00±0.00 | 0.90±0.03 | 0.99±0.00 | 0.87±0.03 | 0.94±0.02 |
| Plasma Cell | L4 | 0.99±0.00 | 0.98±0.01 | 1.00±0.00 | 0.98±0.01 | 0.97±0.02 |
| Myeloid Blast | M1 | 0.98±0.00 | 0.73±0.03 | 0.95±0.01 | 0.69±0.07 | 0.78±0.04 |
| Promyelocyte | M2 | 0.99±0.00 | 0.80±0.03 | 0.97±0.00 | 0.78±0.03 | 0.81±0.06 |
| Myelocyte | M3 | 0.98±0.00 | 0.72±0.04 | 0.97±0.00 | 0.75±0.04 | 0.70±0.08 |
| Metamyelocyte | M4 | 1.00±0.00 | 0.87±0.01 | 0.99±0.00 | 0.85±0.01 | 0.89±0.02 |
| Band Neutrophil | M5 | 1.00±0.00 | 0.95±0.01 | 0.99±0.00 | 0.94±0.01 | 0.95±0.01 |
| Segmented Neutrophil | M6 | 1.00±0.00 | 0.97±0.01 | 1.00±0.00 | 0.98±0.00 | 0.96±0.02 |
| Monocyte | MO2 | 0.99±0.00 | 0.83±0.01 | 0.98±0.00 | 0.82±0.04 | 0.85±0.03 |
| Megakaryocyte | PL2 | 1.00±0.00 | 0.85±0.02 | 0.98±0.00 | 0.74±0.03 | 1.00±0.00 |
| Platelet Clump | PL3 | 0.87±0.05 | 0.74±0.04 | 0.98±0.00 | 0.99±0.00 | 0.59±0.06 |
| Artifact | U1 | 1.00±0.00 | 0.81±0.05 | 0.99±0.00 | 0.69±0.07 | 0.98±0.02 |
| Mitotic Body | U4 | 0.97±0.02 | 0.86±0.02 | 0.99±0.00 | 0.92±0.04 | 0.81±0.02 |
| **Mean** |  | 0.98±0.00 | 0.85±0.01 | 0.99±0.00 | 0.86±0.01 | 0.85±0.01 |
| **Median** |  | 0.99±0.00 | 0.85±0.02 | 0.99±0.00 | 0.86±0.02 | 0.88±0.01 |

**Table 3.** DeepHeme vs Experts F1-Score. F1-score for each of five cross-validation (cv) trained versions of the DeepHeme algorithm, as well as each of the individual export hematopathologist annotators.

| Cell Code | DH_cv1 | DH_cv2 | DH_cv3 | DH_cv4 | DH_cv5 | Expert1 | Expert2 | Expert3 |
| --- | --- | --- | --- | --- | --- | --- | --- | --- |
| B1 | 0.89 | 0.92 | 0.89 | 0.92 | 0.91 | 0.76 | 0.92 | 0.84 |
| B2 | 0.81 | 0.87 | 0.82 | 0.87 | 0.86 | 0.75 | 0.87 | 0.74 |
| E1 | 0.94 | 0.98 | 0.96 | 0.96 | 0.96 | 0.81 | 0.89 | 0.85 |
| E4 | 0.92 | 0.98 | 0.96 | 0.94 | 0.96 | 0.96 | 0.90 | 0.94 |
| ER1 | 0.83 | 0.82 | 0.81 | 0.79 | 0.86 | 0.62 | 0.68 | 0.65 |
| ER2 | 0.76 | 0.75 | 0.81 | 0.70 | 0.78 | 0.52 | 0.69 | 0.65 |
| ER3 | 0.91 | 0.88 | 0.87 | 0.91 | 0.85 | 0.40 | 0.69 | 0.59 |
| ER4 | 0.98 | 0.98 | 0.94 | 1.00 | 0.94 | 0.71 | 0.84 | 0.71 |
| ER5 | 0.96 | 0.94 | 0.96 | 0.94 | 0.98 | 0.81 | 0.92 | 0.92 |
| ER6 | 0.96 | 0.96 | 0.96 | 0.96 | 0.98 | 0.86 | 0.96 | 0.87 |
| L2 | 0.90 | 0.90 | 0.94 | 0.92 | 0.94 | 0.80 | 0.91 | 0.77 |
| L4 | 0.96 | 0.92 | 0.96 | 0.92 | 0.98 | 0.77 | 0.96 | 0.89 |
| M1 | 0.71 | 0.67 | 0.78 | 0.72 | 0.75 | 0.50 | 0.63 | 0.53 |
| M2 | 0.84 | 0.79 | 0.87 | 0.86 | 0.83 | 0.50 | 0.64 | 0.44 |
| M3 | 0.82 | 0.83 | 0.85 | 0.83 | 0.84 | 0.53 | 0.61 | 0.57 |
| M4 | 0.90 | 0.84 | 0.87 | 0.80 | 0.90 | 0.26 | 0.60 | 0.64 |
| M5 | 0.85 | 0.91 | 0.81 | 0.78 | 0.83 | 0.69 | 0.73 | 0.71 |
| M6 | 0.88 | 0.98 | 0.83 | 0.92 | 0.87 | 0.81 | 0.84 | 0.80 |
| MO2 | 0.94 | 0.92 | 0.89 | 0.92 | 0.92 | 0.82 | 0.89 | 0.67 |
| PL2 | 1.00 | 1.00 | 0.98 | 0.98 | 0.98 | 0.77 | 1.00 | 1.00 |
| PL3 | 0.96 | 1.00 | 0.98 | 1.00 | 0.98 | 0.96 | 1.00 | 0.96 |
| U1 | 0.90 | 0.91 | 0.92 | 0.92 | 0.96 | 0.67 | 0.92 | 0.76 |
| U4 | 0.98 | 0.89 | 0.89 | 0.94 | 0.94 | 0.86 | 0.92 | 0.81 |
| Mean | 0.90 | 0.90 | 0.89 | 0.89 | 0.90 | 0.70 | 0.83 | 0.75 |
| Median | 0.90 | 0.91 | 0.89 | 0.92 | 0.92 | 0.76 | 0.89 | 0.76 |

**Table 4.** DeepHeme vs Experts Precision. Precision for each of five cross-validation (cv) trained versions of the DeepHeme algorithm, as well as each of the individual export hematopathologist annotators.

| Cell Code | DH_cv1 | DH_cv2 | DH_cv3 | DH_cv4 | DH_cv5 | Expert1 | Expert2 | Expert3 |
| --- | --- | --- | --- | --- | --- | --- | --- | --- |
| B1 | 0.83 | 0.89 | 0.83 | 0.89 | 0.83 | 0.70 | 0.89 | 1.00 |
| B2 | 0.86 | 0.95 | 0.95 | 0.95 | 1.00 | 0.78 | 0.95 | 0.69 |
| E1 | 0.92 | 1.00 | 0.96 | 0.96 | 0.96 | 0.71 | 0.83 | 0.79 |
| E4 | 0.96 | 0.96 | 0.96 | 0.96 | 0.96 | 0.93 | 0.92 | 0.92 |
| ER1 | 0.87 | 0.81 | 1.00 | 0.83 | 0.85 | 0.55 | 0.73 | 0.67 |
| ER2 | 0.94 | 0.78 | 0.78 | 0.83 | 0.79 | 0.52 | 0.67 | 0.60 |
| ER3 | 0.86 | 0.85 | 0.80 | 0.83 | 0.81 | 0.53 | 0.71 | 0.58 |
| ER4 | 0.96 | 1.00 | 1.00 | 1.00 | 0.92 | 0.74 | 0.95 | 0.65 |
| ER5 | 0.96 | 0.96 | 1.00 | 0.96 | 1.00 | 1.00 | 0.96 | 0.92 |
| ER6 | 0.96 | 0.96 | 0.93 | 0.96 | 0.96 | 0.76 | 0.93 | 0.91 |
| L2 | 0.88 | 0.88 | 0.92 | 0.92 | 0.92 | 0.90 | 0.86 | 0.69 |
| L4 | 0.96 | 0.96 | 0.96 | 0.92 | 1.00 | 0.65 | 1.00 | 1.00 |
| M1 | 0.65 | 0.70 | 0.72 | 0.68 | 0.78 | 0.67 | 0.59 | 0.60 |
| M2 | 0.77 | 0.70 | 0.80 | 0.77 | 0.79 | 0.52 | 0.68 | 0.50 |
| M3 | 0.95 | 0.87 | 0.81 | 0.87 | 0.84 | 0.47 | 0.55 | 0.47 |
| M4 | 0.88 | 0.84 | 0.95 | 0.80 | 0.92 | 0.67 | 0.72 | 0.74 |
| M5 | 0.81 | 0.86 | 0.76 | 0.77 | 0.79 | 0.71 | 0.63 | 0.74 |
| M6 | 0.88 | 1.00 | 0.87 | 0.92 | 0.91 | 0.76 | 0.84 | 0.73 |
| MO2 | 0.96 | 0.96 | 0.95 | 0.92 | 0.92 | 0.83 | 0.95 | 0.82 |
| PL2 | 1.00 | 1.00 | 0.96 | 0.96 | 0.96 | 0.89 | 1.00 | 1.00 |
| PL3 | 1.00 | 1.00 | 1.00 | 1.00 | 1.00 | 1.00 | 1.00 | 1.00 |
| U1 | 0.88 | 0.86 | 0.89 | 0.89 | 0.96 | 0.52 | 0.92 | 0.70 |
| U4 | 1.00 | 0.95 | 0.95 | 1.00 | 1.00 | 1.00 | 0.96 | 1.00 |
| Mean | 0.90 | 0.90 | 0.90 | 0.90 | 0.91 | 0.73 | 0.84 | 0.77 |
| Median | 0.92 | 0.95 | 0.95 | 0.92 | 0.92 | 0.71 | 0.89 | 0.74 |

**Table 5.** DeepHeme vs Experts Recall. Recall for each of five cross-validation (cv) trained versions of the DeepHeme algorithm, as well as each of the individual export hematopathologist annotators.

| Cell Code | DH_cv1 | DH_cv2 | DH_cv3 | DH_cv4 | DH_cv5 | Expert1 | Expert2 | Expert3 |
| --- | --- | --- | --- | --- | --- | --- | --- | --- |
| B1 | 0.96 | 0.96 | 0.96 | 0.96 | 1.00 | 0.84 | 0.96 | 0.72 |
| B2 | 0.76 | 0.80 | 0.72 | 0.80 | 0.76 | 0.72 | 0.80 | 0.80 |
| E1 | 0.96 | 0.96 | 0.96 | 0.96 | 0.96 | 0.96 | 0.96 | 0.92 |
| E4 | 0.88 | 1.00 | 0.96 | 0.92 | 0.96 | 1.00 | 0.88 | 0.96 |
| ER1 | 0.80 | 0.84 | 0.68 | 0.76 | 0.88 | 0.72 | 0.64 | 0.64 |
| ER2 | 0.64 | 0.72 | 0.84 | 0.60 | 0.76 | 0.52 | 0.72 | 0.72 |
| ER3 | 0.96 | 0.92 | 0.96 | 1.00 | 0.88 | 0.32 | 0.68 | 0.60 |
| ER4 | 1.00 | 0.96 | 0.88 | 1.00 | 0.96 | 0.68 | 0.76 | 0.80 |
| ER5 | 0.96 | 0.92 | 0.92 | 0.92 | 0.96 | 0.68 | 0.88 | 0.92 |
| ER6 | 0.96 | 0.96 | 1.00 | 0.96 | 1.00 | 1.00 | 1.00 | 0.84 |
| L2 | 0.92 | 0.92 | 0.96 | 0.92 | 0.96 | 0.72 | 0.96 | 0.88 |
| L4 | 0.96 | 0.88 | 0.96 | 0.92 | 0.96 | 0.96 | 0.92 | 0.80 |
| M1 | 0.80 | 0.64 | 0.84 | 0.76 | 0.72 | 0.40 | 0.68 | 0.48 |
| M2 | 0.92 | 0.92 | 0.96 | 0.96 | 0.88 | 0.48 | 0.60 | 0.40 |
| M3 | 0.72 | 0.80 | 0.88 | 0.80 | 0.84 | 0.60 | 0.68 | 0.72 |
| M4 | 0.92 | 0.84 | 0.80 | 0.80 | 0.88 | 0.16 | 0.52 | 0.56 |
| M5 | 0.88 | 0.96 | 0.88 | 0.80 | 0.88 | 0.68 | 0.88 | 0.68 |
| M6 | 0.88 | 0.96 | 0.80 | 0.92 | 0.84 | 0.88 | 0.84 | 0.88 |
| MO2 | 0.92 | 0.88 | 0.84 | 0.92 | 0.92 | 0.80 | 0.84 | 0.56 |
| PL2 | 1.00 | 1.00 | 1.00 | 1.00 | 1.00 | 0.68 | 1.00 | 1.00 |
| PL3 | 0.92 | 1.00 | 0.96 | 1.00 | 0.96 | 0.92 | 1.00 | 0.92 |
| U1 | 0.92 | 0.96 | 0.96 | 0.96 | 0.96 | 0.92 | 0.92 | 0.84 |
| U4 | 0.96 | 0.84 | 0.84 | 0.88 | 0.88 | 0.76 | 0.88 | 0.68 |
| Mean | 0.90 | 0.90 | 0.89 | 0.89 | 0.90 | 0.71 | 0.83 | 0.75 |
| Median | 0.92 | 0.92 | 0.92 | 0.92 | 0.92 | 0.72 | 0.88 | 0.80 |

**Figure 1.**
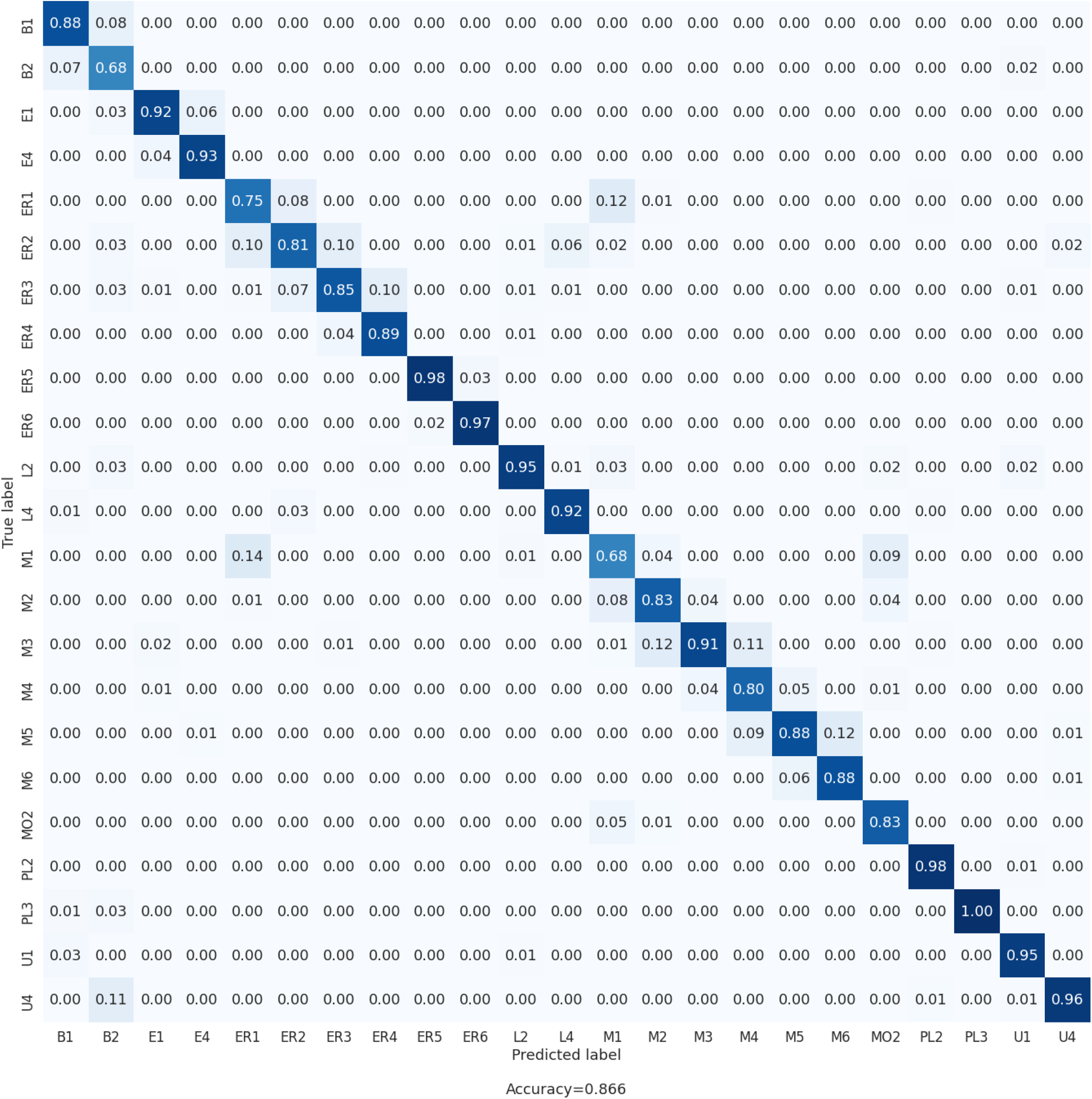
Confusion Matrix. This figure shows the confusion matrix of prediction on the test set of UCSF images. Most misclassifications are between biologically adjacent cell classes, reflecting the true ambiguity between edge cases. Notable examples include myeloid blast (M1) vs erythroid blast (ER1). These are developmentally adjacent cell types and as a result have some morphologic overlap.

**Figure 2.**
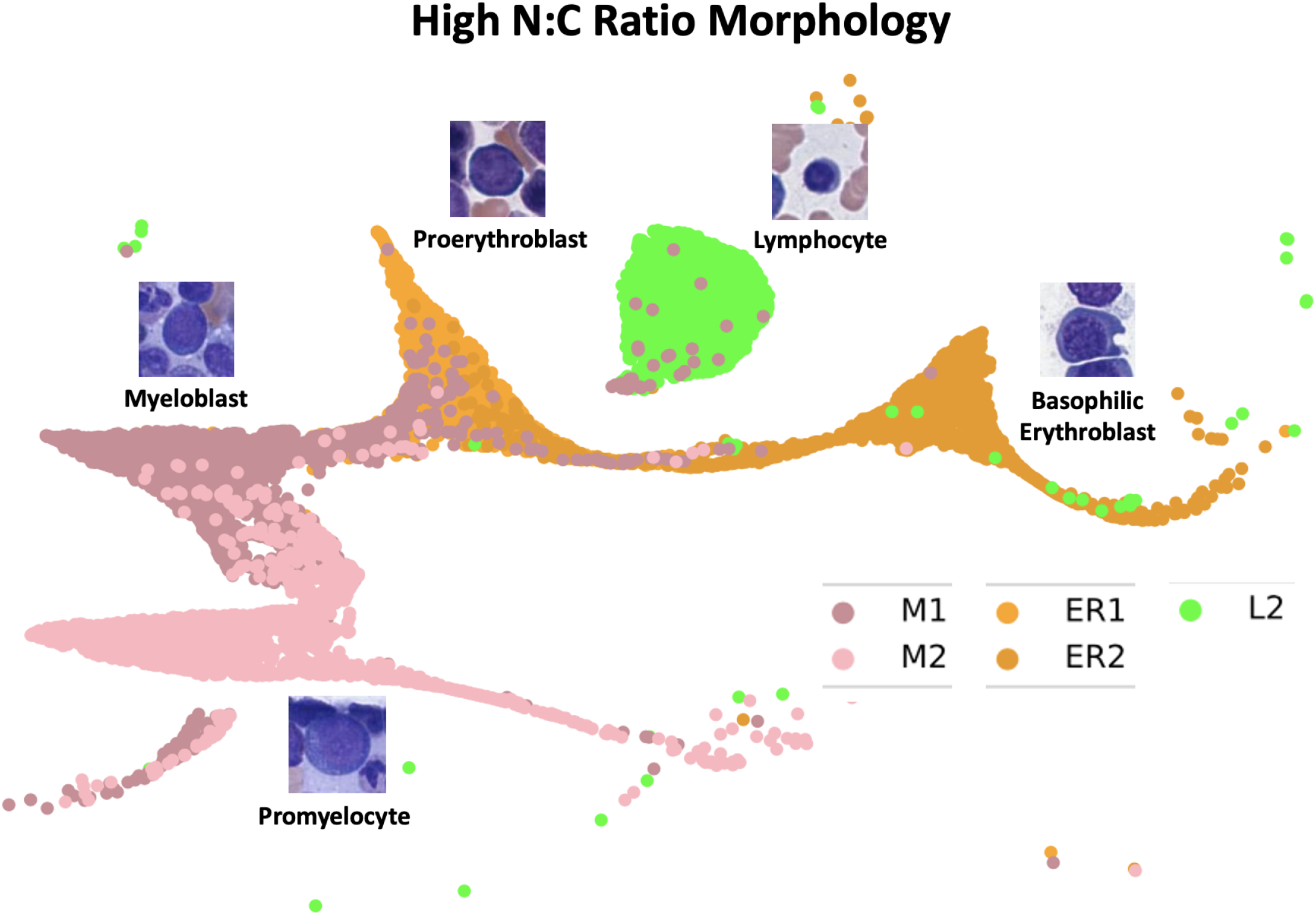
UMAP embedding for cells with high nucleus:cytoplasm (N:C) ratio. All five cell classes in our dataset with high N:C ratio co-localize, while only cell clusters that are directly related to each other are attached by bridges. Of note, the bridge between myeloblasts (M1) and proerythroblasts (ER1) reflects the location of the theoretical hematopoietic stem cell that exists as a precursor between them.

**Figure 3.**
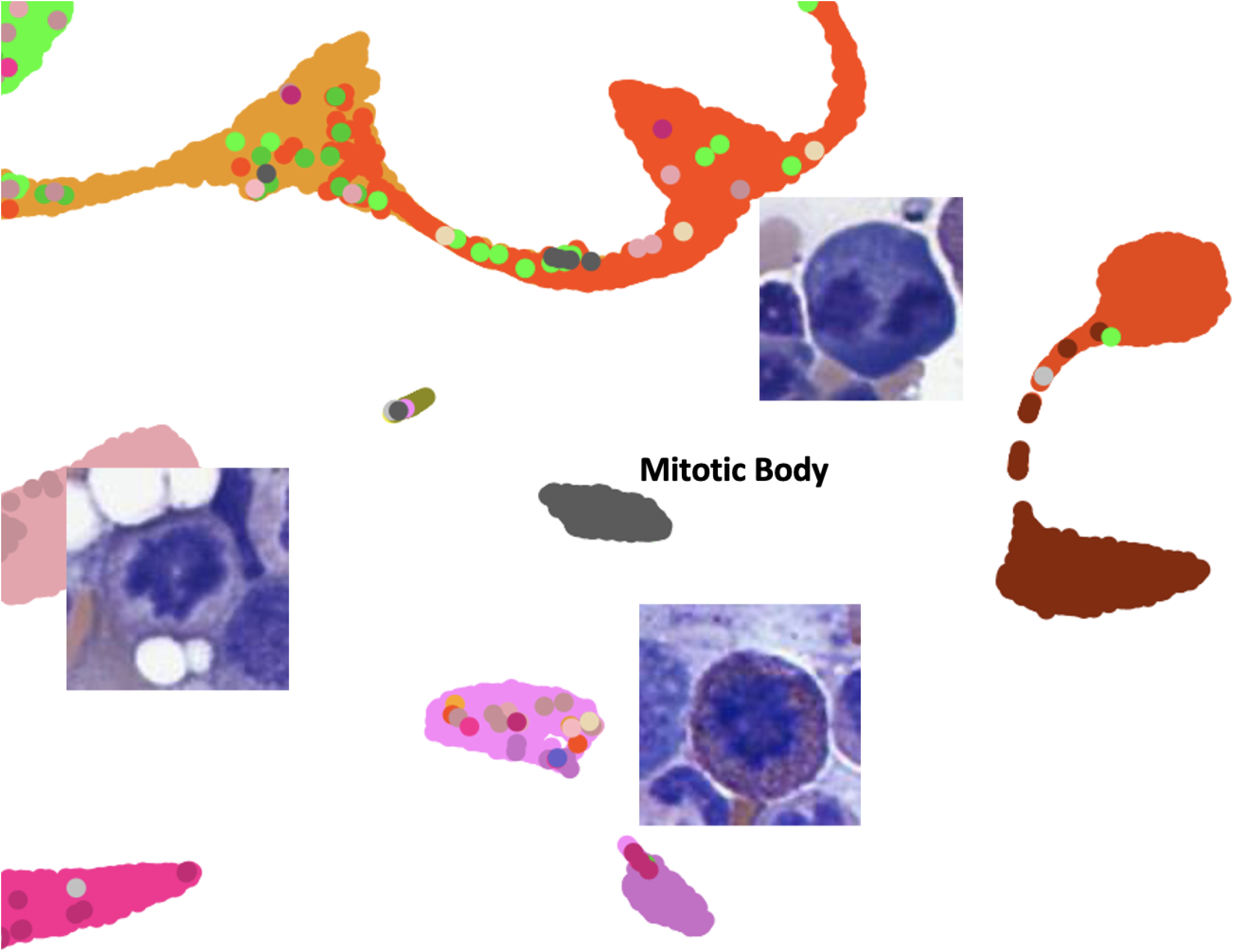
UMAP embedding for mitotic bodies. The mitotic bodies in this study broadly derive from three morphologic groups based on cytoplasmic color, which correspond to the lineage of the cell undergoing mitosis. Mitotic cells in the neutrophil lineage have a pale pink cytoplasm, those in the eosinophilic lineage contain bright pink cytoplasm and those in the erythroid lineage have deep blue cytoplasm. The mitotic body cluster is placed by the UMAP directly between these three lineage cluster sets, perhaps reflecting the relationship of the mitotic body class to these three lineages. Further subclassification of mitotic bodies by lineage is possible based on cytoplasm color.

## Notes

### Competing Interest Statement

The authors have declared no competing interest.

